# Predicted sensory consequences of voluntary actions modulate amplitude and temporal dynamics of preceding readiness potentials

**DOI:** 10.1101/101402

**Authors:** Daniel Reznik, Shiri Simon, Roy Mukamel

## Abstract

Self-generated, voluntary actions, are preceded by a slow negativity in the scalp electroencephalography (EEG) signal recorded from frontal regions (termed ‘readiness potential’; RP). This signal, and its lateralized subcomponent (LRP), is mainly regarded as preparatory motor activity associated with the forthcoming motor act. However, it is not clear whether this neural signature is associated with preparatory motor activity, expectation of its associated sensory consequences, or both. Here we recorded EEG data from 12 healthy subjects while they performed self-paced button presses with their right index and middle fingers. In one condition (motor+sound) these button-presses triggered a sound while in another (motor-only) they did not. Additionally, subjects passively listened to sounds delivered in expected timings (sound-only). We found that the RP amplitude (locked to time of button press) was significantly more negative in the motor+sound compared with motor-only conditions starting ~1.4 seconds prior to button press. Importantly, no signal negativity was observed prior to expected sound delivery in the sound-only condition. Thus, the differences in RP amplitude between motor+sound and motor-only conditions are beyond differences in mere expectation of a forthcoming auditory sound. No significant differences between the two conditions were obtained in the LRP component. Our results suggest that expected auditory consequences are encoded in the early phase of the RP preceding the voluntary actions that generate them.

## Introduction

Self-generated, voluntary actions are rarely performed without a preceding preparation or planning period during which various decisions regarding the upcoming actions’ time, trajectory and goal are made. Accumulating research over the past decades suggests that neural activity during the time interval preceding voluntary action execution is associated with different aspects of the forthcoming motor act, such as task, action type, and selection of appropriate effector (for a review see Haggard, 2008). This view is supported by functional magnetic resonance imaging (fMRI) and electrophysiological studies in humans and primates showing that the neural activity during preparatory time intervals preceding action execution can represent, for example, the type of actions (grasping or touching; Gallivan, McLean, Smith, & Culham, 2011), the executing effector (right or left hand; Soon, Brass, Heinze, & Haynes, 2008) or the tool that is about to be used (Brandi, Wohlschlager, Sorg, & Hermsdorfer, 2014; see also Bulea, Prasad, Kilicarslan, & Contreras-Vidal, 2014; Cisek & Kalaska, 2004; Perez et al., 2015)

Self-initiated, voluntary actions are usually preceded by a slow negativity in the scalp electroencephalography (EEG) recorded from frontal and central regions (termed “readiness potential”, RP; Kornhuber & Deecke, 1990; Libet, Gleason, Wright, & Pearl, 1983). This negativity, is usually divided into early and late phases, starting around ~1500 ms and ~500 ms prior to action execution, respectively (Shibasaki & Hallett, 2006). The early phase is believed to reflect gradual increase in neural firing rate in high motor cortical regions associated with initiatory and preparatory motor activity, such as supplementary motor area (SMA; Lang et al., 1991; Yazawa et al., 2000; Fried, Mukamel, & Kreiman, 2011; Pedersen et al., 1998; Cunnington, Windischberger, Deecke, & Moser, 2003). The late phase is more lateralized and specific to the motor command of the executing effector and is therefore believed to represent preparatory activity in primary motor cortex (M1; Passingham, 1987; Pedersen et al., 1998).

When unimanual voluntary action is performed, for example by a right hand, stronger RP negativity is usually found over the left hemisphere - contralateral to the active hand (Kutas & Donchin, 1980). This difference in RP has been shown to begin in close temporal proximity to initiation of the motor act (De Jong, Wierda, Mulder, & Mulder, 1988; Gratton, Coles, Sirevaag, Eriksen, & Donchin, 1988; Smid, Mulder, & Mulder, 1987) and is usually detected by subtracting the RP signal over the ipsilateral hemisphere from that of the contralateral hemisphere with respect to the active hand. This late RP component, termed the lateralized readiness potential (LRP) is interpreted as manifestation of preparatory brain activity that is more specific to the executing effector which is about to perform the action.

Despite recent evidence suggesting that RP does not necessarily reflect motor preparation, but might be associated with general decision making processes (Alexander et al., 2016), this neural signature is predominantly discussed in the framework of a neural correlate of motor intention and consciousness of such intention (Haggard & Eimer, 1999; Libet et al., 1983). Accordingly, only few studies examined the relation of RP to self-generated sensory consequences (Ford, Palzes, Roach, & Mathalon, 2014) and their temporal perception (Jo, Wittmann, Hinterberger, & Schmidt, 2014). The link between RP and self-generated sensory consequences can be formulated within the forward model of sensory-motor-integration (Wolpert, Ghahramani, & Jordan, 1995). According to the model, during the preparation period prior to voluntary action execution, the motor cortex sends signals both to the executing effectors and to the relevant sensory cortex, in which the sensory consequences of the action are expected. The signal that is sent from motor to sensory cortices was termed “efference copy” (von Holst, 1954) and was proposed to facilitate, attenuate, or otherwise modulate both perceptual and neural responses to self-generated stimuli compared with responses to identical stimuli generated by an external source (see Crapse & Sommer, 2008 for a review). Therefore, the RP and its LRP subcomponent, are candidate neurophysiological signatures for such a forward model embedding the expected sensory consequences of self-generated actions.

The aim of the current study was to examine whether the RP and\or LRP encode the expected sensory consequences triggered by voluntary actions and characterize the temporal aspects of such encoding. To this end, we recorded EEG data from healthy subjects while they performed voluntary, self-paced button presses. We found that when button presses were associated with auditory consequences, the RP preceding such presses was more negative compared with RPs preceding identical button presses that were not associated with auditory consequences. Our results suggest that the motor activity preceding voluntary actions, as reflected by the early phase of EEG recorded RP, encodes the expected auditory consequences and that such encoding in turn can modulate the subsequent sensory-evoked activity in auditory cortex.

## Experimental methods

### Subjects

Seventeen healthy, right-handed undergraduate students naïve to the purposes of the study were recruited to the experiment. The data from three subjects were excluded due to technical problems during EEG recording, thus leaving data from 14 subjects (four males; mean age: 23, rage: 19 - 26 years) for subsequent analysis. The study conformed to the guidelines that were approved by the ethical committee in Tel-Aviv University. All subjects provided written informed consent to participate in the study and were compensated for their time.

### Procedure

Subjects were seated in a dimly lit room and performed self-paced button presses with index or middle fingers of their right hand while fixating on a cross (“+”) constantly displayed on a computer screen. The task consisted of two consecutive runs - in one run, middle finger button-presses triggered an auditory stimulus (300 ms C major piano chord generated by MIDI-OX software ver. 7.0.2, delivered through free field speakers; motor+sound condition), while index finger button-presses did not trigger any auditory stimulus (motor-only condition); in the other run, the finger mapping with respect to sound was reversed (middle finger button press = motor-only condition; index finger button press = motor+sound condition). The order of runs (finger-mapping) was counterbalanced across subjects. Before the experimental procedure, subjects were informed about the finger-sound mapping and were instructed to perform the button presses in a self-paced manner (with at least 3-4 seconds between consecutive presses), freely choosing between index and middle finger. If inter press interval (IPI) of two consecutive presses was less than 3.5 seconds, the fixation point changed its color to red for 500 ms to inform the subject of insufficient IPI and the data related to the last button-press were discarded from analysis. Each run ended when subjects performed at least 70 “good” presses (with IPI greater than 3.5 seconds) with each finger.

**Figure 1.**
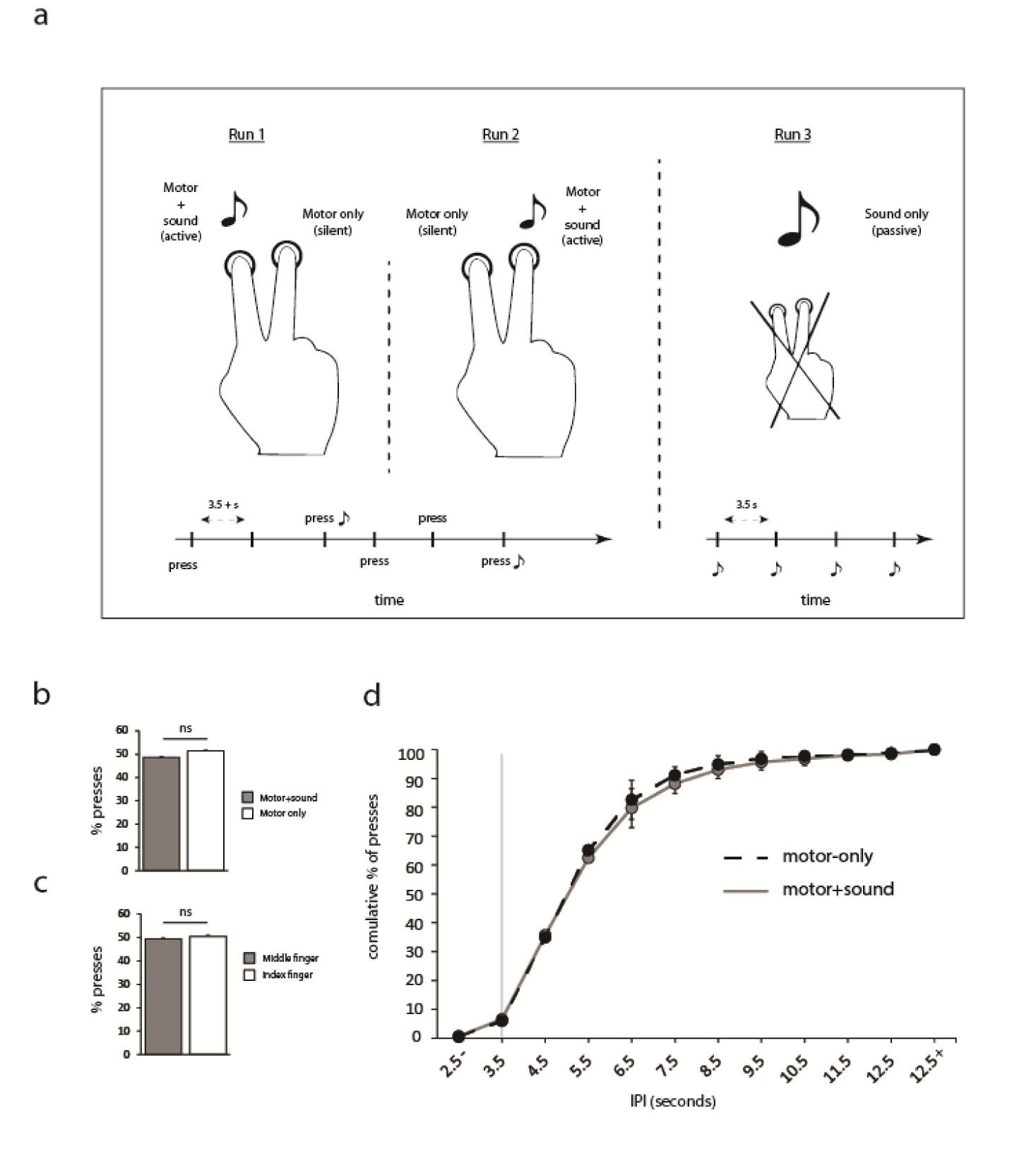
**Experimental design and behavioral results.** (**a**) The experiment consisted of three runs. In the first two runs, subjects performed voluntary self-paced button presses with their index and middle fingers. In one run, index finger presses triggered auditory stimuli (motor+sound condition), while middle finder presses did not (motor-only condition; Run 1 in the Figure). In the other run, the finger mapping was reversed, such that middle finger presses triggered auditory stimuli and index finger presses did not (Run 2 in the Figure). After the two runs, subjects passively perceived the auditory stimuli delivered at a fixed rate (Run 3 in the Figure). (**b**) Mean±s.e.m. percentage of presses across subjects was not different between motor+sound and motor-only conditions (paired t-test, n.s.). (**c**) Likewise, the mean±s.e.m. percentage of index and middle finger presses was not significantly different across subjects. (**d**) Cumulative distribution of inter-press-intervals across subjects in motor+sound and motor-only condition was not significantly different. The vertical line denotes lowest IPI limit for definition of “good” trials taken for analysis. Each data point denotes mean±st.d. across subjects, percentage of presses with lower IPI (total of 12 time bins; repeated measures ANOVA, n.s.).

To allow us to distinguish effects of brain activity evoked by mere expectation of auditory stimuli (without a motor component), after the two active runs subjects underwent an additional run, during which they passively listened to 70 repetitions of the same chord presented in the motor+sound condition, delivered in a fixed rate of once every 3.5 seconds (sound-only condition; Figure 1a). In order to keep subjects engaged during this run, they performed an oddball detection task (300 ms, 500 Hz pure tone; additional ten pseudo-randomly interspersed odd-ball trials per run). Detection of such oddballs was reported by a button press and data from these catch trials was not analyzed.

### EEG recording and data analysis

EEG signals were recorded using a BIOSEMI Active II 64-channel system at a sampling rate of 256 Hz. Additional 3 channels were used to record activity related to ocular artifacts and 2 more channels placed on right and left mastoids served as offline reference channels.

The EEG data was analyzed using Matlab (2013b; MathWorks) and EEGLAB toolbox (version 12.0.1). The continuous data was re-referenced to the average of the mastoid channels and off-line filtered with a window based finite impulse response 0.05-40 Hz band pass filter implemented in EEGLAB toolbox. We segmented the data into epochs covering the time window from −2.6 to 0.6 seconds relative to trial onset (time 0; a button press in the motor+sound and motor-only conditions, and sound delivery in the sound-only condition). For removing ocular artifacts, we used independent component analysis (ICA) implemented in EEGLAB toolbox. Ocular ICA components (range across subjects: 1 to 2 components) were identified by visual inspection and deleted from the global signal. Noisy trial epochs (exceeding ±100 µV range) at Fz, FCz, Cz, C3 and C4 channels were identified in the raw data and rejected from the analysis. Finally, for each subject the number of trials in motor-only, motor+sound and sound-only conditions was balanced by taking the first number of trials matching the condition with the lowest number of trials remaining after pre-processing.

Pre-stimulus event related potentials (ERPs) were analyzed at Fz, FCz, Cz, C3 and C4 channels using a baseline period from −2600 ms to −2500 ms (Haggard & Eimer, 1999), relative to event onset. Lateralized readiness potential (LRP; Gratton, Coles, Sirevaag, Eriksen, & Donchin, 1988) was measured by calculating the difference between readiness potentials recorded from C3 and C4 channels (C3 - C4). For estimating RP and LRP amplitudes, for each subject we calculated the mean signal during the time window −2500 to 0 ms prior to button press.

Differences in the RP and LRP between motor+sound and motor-only conditions could arise from audio-motor coupling encoding the auditory consequences of self-generated actions, but also merely from differences in sensory anticipation present in the motor+sound and absent in the motor-only condition. To address this issue we applied a commonly used procedure for correcting the EEG signal preceding motor+sound trials by subtracting from it the EEG activity preceding sound-only trials (e.g., Baess et al., 2011; Horvath, Maess, Baess, & Toth, 2012). Since the timing of auditory sounds in the sound-only condition was expected, differences between the corrected motor+sound EEG signal and sound-only EEG signal should reflect differences that are not related to differences in expectancy of an upcoming auditory stimulus but more plausibly reflect differences in sensory-motor coupling. Similarly, LRP were corrected by subtracting the difference between C3 and C4 channels in sound-only trials from the same difference in the motor+sound trials.

We also examined potential effects of the RP on the EEG signal following sound onset. For assessing the evoked N1 component, we used a 100 ms baseline prior to sound delivery (in sound-only and motor+sound conditions). We first calculated the averaged evoked responses across subjects and chanels (Fz, FCz and Cz) separately for motor+sound and sound-only conditions and determined the time to peak of the N1 component in each condition. In accordance with previous studies (Baess, Horvath, Jacobsen, & Schroger, 2011), to calculate the N1 amplitude in individual subjects, we calculated the average signal in a 43 ms window centered around the group peak time in the corresponding condition. Additionally, to correct for the motor component involved in N1 amplitude evoked during motor+sound condition (and absent in the sound-only condition), we subtracted the evoked activity during motor-only trials from the motor+sound condition before comparing it with the evoked response of the sound-only condition. All statistical comparisons were conducted using significance level of α = 0.05.

## Results

Two subjects were excluded from the analysis since they did not exhibit a negative trend in voltage (readiness potential) prior to movement onset both in motor+sound and motor-only conditions. Similar exclusion rates due to lack of RP have been previously reported (Schurger, Sitt, & Dehaene, 2012). All further analyses were conducted on the remaining 12 subjects.

At the behavioral level, there was no significant difference between the proportion of times subjects performed button presses that were associated or not with a sound (motor+sound: 49.40±0.91%, motor-only: 50.60±0.91%; t_(23)_=0.63, p=0.534; Figure 1b), or the proportion of times subjects pressed using their index/middle finger (index finger: 51.30±0.88%, middle finger: 48.70±0.88%; mean±s.e.m across subjects; two-tailed paired t-test, t_(23)_=1.51, p=0.143; Figure 1c). Next, we calculated the distribution of inter-press-intervals for the motor+sound and motor-only conditions. We found no significant difference in the distribution of inter-press-intervals (IPI; binned into 12 time intervals) across subjects for the two conditions (repeated measures analysis of variance – condition X time bins, Greenhouse-Geisser corrected F_(3,41)_=1.327, p=0.277; Figure 1d). The mean IPI across conditions and subjects was 5.47±0.28 seconds, with larger mean IPI in the motor+sound compared with motor-only condition (motor+sound: 5.56±0.30 seconds, motor-only: 5.39±0.26 seconds, two-tailed paired t-test, t_(11)_=2.29, p=0.042). The mean percentage of discarded trials (with IPI less than 3.5 seconds) across subjects was 6.2% with no significant difference between motor+sound and motor-only conditions (motor+sound: 6.49±0.92%, motor-only: 6.02±0.89%, two-tailed paired t-test, t_(11)_=0.73, p=0.481). The mean number of balanced trials across subjects that were analyzed after preprocessing was 66 (range: 62-70 trials). Subjects had perfect performance in the odd-ball trials detection task.

At the physiological level we first examined the differences in the mean RP amplitude across Fz, FCz, Cz channels during the epoch preceding button presses in motor+sound and motor-only conditions. We found that the mean RP amplitude was more negative before motor+sound compared with motor-only conditions (mean±s.e.m µV across subjects – motor+sound condition: −1.69±0.30 µV, motor-only condition: −0.68±0.35 µV, paired one-tailed t-test, t_(11)_=2.36, p=0.018; Cohen’s d=0.71). Importantly, after controlling for auditory expectancy during motor+sound condition (see methods for details), we found that the mean RP across Fz, FCz and Cz channels was still more negative in the motor+sound compared with motor-only conditions (mean±s.e.m µV across subjects – motor+sound_corr_ condition: −1.83±0.44 µV, paired one-tailed t-test, t_(11)_=2.12, p=0.028; Cohen’s d=0.63). Thus, the stronger negativity preceding the motor+sound compared with motor-only conditions is unlikely to be explained by mere differences in expectancy of a forthcoming auditory stimulus devoid of a motor component.

**Figure 2.**
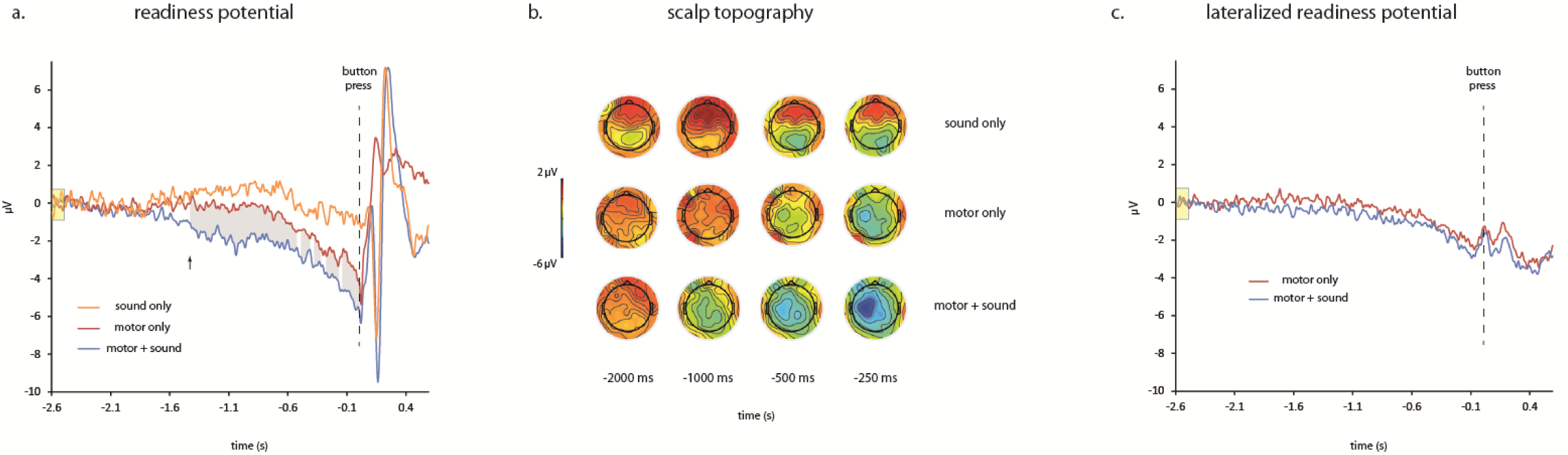
**Readiness Potentals.** (**a**) Mean readiness potential across subjects was significantly more negative in the motor+sound condition compared with the motor-only condition starting −1.43 seconds (marked with an arrow) relative to time of button press. The shaded area denotes significant difference (corrected for q(FDR)<0.05) between the two readiness potential time-courses. The signal preceding sound delivery in sound-only condition was not significantly different than zero at any time point. The marked yellow area denotes the time interval used as baseline (**b**) Group (n=12) topographical maps showing scalp activity prior to button presses in motor+sound and motor-only conditions and sound delivery in sound-only condition. Note the early left-oriented negativity during the time preceding sound triggering button presses. (**c**) Lateralized readiness potential time-courses. The mean LRP of the entire time window prior to button press [-2500:0] of the motor+sound condition was significantly more negative than the motor-only condition but not at any individual time point. The marked yellow area denotes the time interval used as baseline

Next, we compared the temporal profile of RP preceding motor-only and motor+sound conditions. To this end, for each subject we calculated the mean RP across trials, collapsed across Fz, FCz, and Cz channels separately for each condition (motor+sound and motor-only). In each time point (defined by EEG sampling rate – 256 Hz), we performed a two-tailed paired t-test across the two conditions, corrected for false discovery rate (FDR) with q(FDR)<0.05 (Benjamini & Hochberg, 1995). This allowed us to calculate time points where the RPs were significantly different. The first significant time point that was followed by at least 26 consecutive significant samples (~100 ms) was considered as the time point were the RPs started to differ. We found that motor+sound RP was significantly more negative than motor-only RP starting 1.43 seconds prior to button press (see group average in Figure 2a and group topological maps in Figure 2b). From this time-point onward until the time of button press, the difference between the RP of the two conditions was significant except for four brief periods not exceeding ~40 ms each. The signal time course preceding the sound-only condition (in which the sound is temporally expected) was not significantly lower than zero at any time point. Accordingly, after controlling for auditory expectancy during the motor+sound condition, we found that the RPs preceding motor+sound and motor-only conditions were still significantly different for a consecutive time window between 1.43 to 0.5 s prior to button press (Figure 3a).

**Figure 3.**
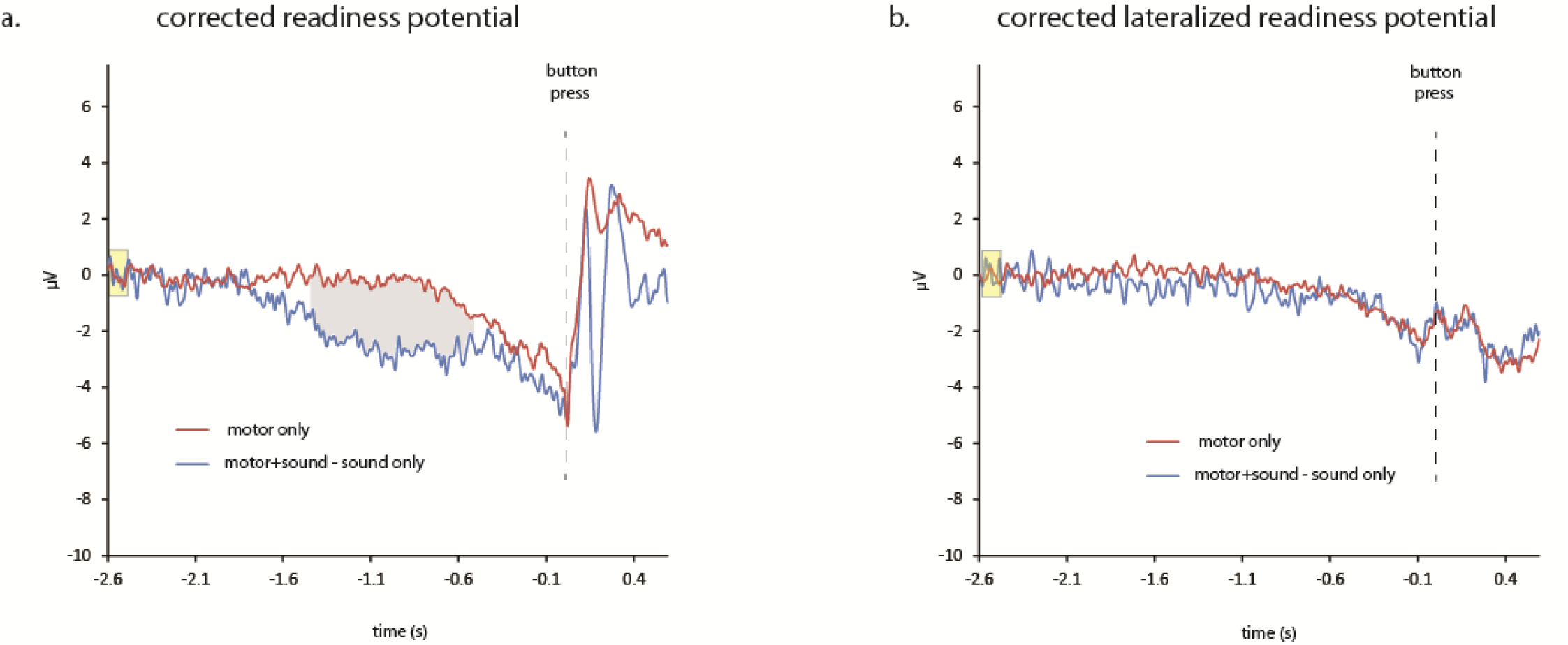
**Readiness potentials corrected for auditory expectation**. **(a)** Mean readiness potentials across subjects after substracting the activity preceeding sound-only condition from the activity preceding motor+sound condition. The shaded area denotes significant difference (corrected for q(FDR)<0.05) between the two readiness potential time-courses. The marked yellow area denotes the time interval used as baseline. (b) Lateralized readiness potential time-courses were not different between the corrected motor+sound and motor-only conditions. The marked yellow area denotes the time interval used as baseline.

To examine the involvement of specific RP components in coding expected auditory action consequences, we also examined the lateralized readiness potentials (LRP; see methods) preceding motor+sound and motor-only conditions. Similar to what we found for the RP, mean LRP amplitude preceding motor+sound condition was more negative compared with LRP amplitude preceding motor-only condition (mean±s.e.m µV across subjects – motor+sound condition: −0.77±0.14 µV, motor-only condition: −0.36±0.14 µV, paired one-tailed t-test, t_(11)_=2.95, p=0.006, Cohen’s d=0.88). However, when we controlled for auditory expectation (see methods for details), the difference was no longer significant (mean±s.e.m µV across subjects – motor+sound_corr_ condition: −0.62±0.28 µV, paired one-tailed t-test, t_(11)_=1.48, p=0.164). When we examined the temporal profile of LRP preceding motor+sound and motor-only conditions, we found that the LRPs were not significantly different between the two conditions at any time point neither before or after correction for auditory expectancy (Figure 2c and Figure 3b; before the correction there was only one significant time point, – 1.47 seconds prior to button press). This suggests that the LRP component is involved to a lesser extent than the RP in encoding auditory consequences of self-generated actions.

Sounds are usually followed by an evoked negativity over scalp EEG midline channels around 100 ms after the sound onset (specifically, N1 component recorded from channels FCz, Fz and Cz). Studies examining how N1 amplitude is modulated by self- compared with externally-generated sounds report attenuated N1 amplitude for self-initiated sounds (Baess et al., 2011; Janos Horvath, Maess, Baess, & Toth, 2012; Hughes, Desantis, & Waszak, 2013). This modulation is believed to occur via “efference copies” (von Holst, 1954) sent from motor to auditory areas for matching between the predicted and actual self-generated action consequences (Wolpert, Ghahramani, & Jordan, 1995). At the temporal domain, the time-to-peak of the mean N1 across subjects and channels (FCz, Fz and Cz) was at 175 ms in the motor+sound condition and 164 ms in the sound-only condition, with no significant difference across subjects (paired two-tailed t-test, t_(11)_=0.62, p=0.545).

We have shown that RP is significantly more negative in the motor+sound (active) condition than in the sound-only (passive) condition in the time interval preceding the button-press/ sound delivery. Since such pre-stimulus differences in baseline can affect the magnitude of the ensuing stimulus-evoked N1 response, we examined this issue using either a baseline of 100 ms prior to sound onset or the baseline we used for RP calculations (-2600 to −2500). We found that the N1 amplitude in the motor+sound condition depended dramatically on the baseline used. Using a baseline of 100 ms prior to sound onset, the N1 amplitude was attenuated in motor+sound compared with sound-only conditions (sound-only: −5.11±0.89 µV, motor+sound: −2.98±0.84 µV, paired two-tailed t-test, t_(11)_=2.19, p=0.050, Cohen’s d=0.61; Figure 4a). When using a baseline of −2600 to −2500 ms prior to sound onset, no such difference in N1 amplitudes was found (sound-only: −5.89±0.83 µV, motor+sound: −8.07±1.26 µV, paired two-tailed t-test, t_(11)_=1.59, p=0.13; Figure 4b). This was mainly due to the changes in N1 amplitude in the motor+sound condition when calculated using the different baselines (mean±s.e.m µV across subjects – [-100 to 0 ms]: −2.98±0.88 µV, [−2600 to −2500 ms]: −8.07±1.26 µV, paired two-tailed t-test, t_(11)_=6.75, p<10^-4^, Cohen’s d=2.15). Conversely, the N1 amplitude in the sound-only condition was invariant to baseline used ([-100 to 0 ms]: −5.11±0.89 µV, [−2600 to −2500 ms]: −5.89±0.83 µV, paired two-tailed t-test, t_(11)_=1.56, p=0.14).

**Figure 4.**
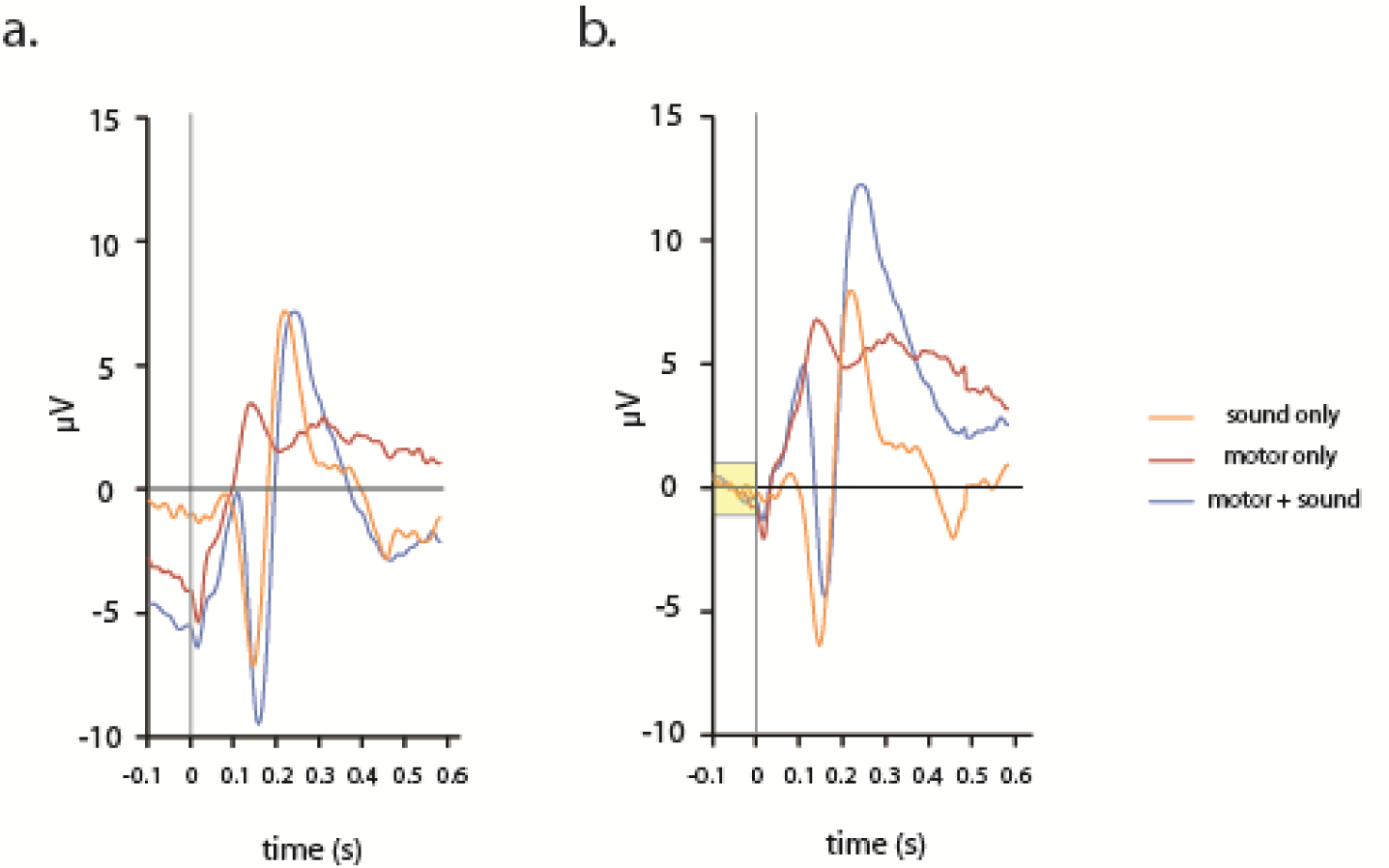
**Evoked auditory N1 component.** (**a**) Using a baseline of 2600 to2500 ms prior to sound onset, the mean N1 amplitude across subjects was not significanly different between motor+sound and sound-only conditions. Note the strong differences in signal during the time-interval of 100 ms prior to event onset which is typically used for N1 baseline calculation. (**b**) Using the common window of 100 ms prior to sound onset as baselin, the N1 amplitude in motor+sound condition was attenuated compared with N1 amplitude in sound-only condition (paired t-test, p<0.05). The marked yellow area denotes the time interval used as baseline. The motor+sound signals presented in the figure are before correction for motor-only component. Correction for the motor-only component resulted in enhancement of motor+sound N1 using both time windows for baseline definitions.

After correcting the N1 amplitude in the motor+sound condition for the activity evoked during motor-only conditions, we found that using either baselines, the corrected N1 amplitude in motor+sound condition was more negative compared with the N1 amplitude in the sound-only condition ([-100 to 0 ms]: sound-only: −4.96±0.91 µV, motor+sound_corr_: −9.11±1.10 µV, one-tailed paired t-test, t_(11)_=3.87, p=0.001, Cohen’s d=1.13; [−2600 to −2500 ms]: sound-only: −5.81±0.86 µV, motor+sound_corr_: −11.00±1.28 µV, t_(11)_=3.95, p=0.001, Cohen’s d=1.10).

## Discussion

In the current study, subjects performed voluntary, self-paced button presses that were either associated with auditory consequences or not. We found that readiness potential prior to sound-triggering button presses was characterized by stronger negativity and earlier onset compared with the readiness potential preceding button presses with no auditory consequences. Importantly, these differences could not be attributed merely to the expectation of an auditory event which is present in one motor act and not the other. Furthermore, we found that, unlike the readiness potential (RP), the amplitude of the lateralized readiness potential (LRP) was not sensitive to differences in the actions’ auditory consequences. Finally, we found that the magnitude of evoked N1 component during perception of self-generated sounds was sensitive to the time window used as baseline.

It has been suggested that sensory regions undergo top-down modulation by the motor system during voluntary action execution (for a review see Crapse & Sommer, 2008; Hughes, Desantis, & Waszak, 2012). Accumulating theoretical (Schurger, Sitt, & Dehaene, 2012; Wolpert et al., 1995) and empirical (Eliades & Wang, 2003; Haggard, Clark, & Kalogeras, 2002; Voss, Ingram, Haggard, & Wolpert, 2006) evidence suggest that the modulatory effect of self-generated actions on neural activity and perception starts *prior* to execution of an action or actual delivery of the re-afferent stimuli. This notion is supported by animal and human studies showing increased synchronization between motor and sensory regions upon action execution such as whisker movement in rats (Nicolelis, Baccala, Lin, & Chapin, 1995) or speech onset in healthy humans (Ford, Roach, Faustman, & Mathalon, 2007). This might explain reports of attenuated sensation even in the absence of a movement (Voss et al., 2006) or perception of self-generated auditory consequences as being temporally closer to the sound-triggering actions (intentional binding phenomenon; Haggard et al., 2002). In the current study, we show that expectation of self-generated action consequences starts to modulate neural activity at the early phase of RP, in the time window between ~1.4 to 0.5 seconds prior to sound-triggering action. This, could potentially activate and prepare the motor-sensory network for perception of self-generated stimuli prior to its actual delivery/generation. Accordingly, Jo and colleagues (Jo et al., 2014) explicitly associated between the early RP phase and behavioral reports of temporal perception of ensuing stimuli in the auditory domain. The authors report that subjects with more negative early RP showed greater intentional binding, namely, these subjects perceived self-generated sounds as being closer in time to the sound triggering actions. Together, these results suggest that the early RP component prior to sound-triggering actions is related to perception of self-generated auditory consequences.

Interestingly, pathological populations, such as schizophrenic patients, exhibit reduced modulations in responses to self- compared to other-generated stimuli. For example, schizophrenic patients do not show reduced tickling sensation of self-generated tactile stimulation compared with healthy controls (Blakemore, Smith, Steel, Johnstone, & Frith, 2000) and struggle in differentiating self-generated action consequences from identical consequences generated by others (Frank, 2001; Waters, Woodward, Allen, Aleman, & Sommers, 2012). It is believed that these phenomena are due to a failure in sensorimotor integration (Wolpert et al., 1995) and predictive coding mechanisms (Ford, Roach, Faustman, & Mathalon, 2008; Frith, Blakemore, & Wolpert, 2000). This view is supported by findings that schizophrenic patients, unlike healthy controls, do not exhibit stronger RP negativity prior to sound-triggering actions compared with RP negativity prior to silent actions (Ford et al., 2014) and fail to synchronize between motor and sensory areas prior to speech onset (Ford et al., 2008). These data in pathological populations further support the notion that preparatory motor activity prior to voluntary actions might embed their upcoming sensory consequences.

In terms of functional neuroanatomy, readiness potential and lateralized readiness potential components are believed to reflect neural activity in different brain areas. While the cortical source of readiness potential is believed to reside in frontal, high-order areas of the motor system, such as SMA and pre-SMA (Lang et al., 1991; Shibasaki & Hallett, 2006; Yazawa et al., 2000), the lateralized readiness potential is believed to be related to more specific, low-level motor components originating in M1 contralateral to the executing effector (Eimer, 1998; Haggard & Eimer, 1999). In the current study we found differences in modulation of RP and LRP prior to sound triggering actions. In agreement with a previous study (Ford et al., 2014), we find more negative RP and LRP amplitudes in the motor+sound compared with motor-only conditions in healthy subjects. However, after controlling for auditory expectancy, the LRP amplitudes were no longer different. This suggests that the encoding of sensory consequences for self-generated actions occurs predominantly in high-order motor areas. This notion is compatible with neuroimaging studies that examined the source of sensory modulation in response to self-generated compared with externally-generated stimuli and pointed to high-order motor areas, such as SMA and lateral pre-motor cortex, as the source of such modulation (Christensen et al., 2007; Cui et al., 2014; Ford et al., 2014; Haggard & Whitford, 2004; Makoshi, Kroliczak, & van Donkelaar, 2011; Reznik, Ossmy, & Mukamel, 2015). Nonetheless, other studies using fMRI show that activity in primary motor cortex is also modulated by self-generated action consequences (Eisenberg, Shmuelof, Vaadia, & Zohary, 2011; Reznik et al., 2015), therefore the role of low level motor regions in encoding sensory consequences of voluntary actions remains to be further elucidated.

Additionally, we found that while N1 amplitude evoked by self-generated sounds was sensitive to the time interval used as a baseline, N1 amplitude evoked by passively perceived sounds was not. As we show, EEG signal dynamics are dramatically different prior to self- and externally generated sounds (Figure 2a, Figure 4a). Therefore, using a time window which is close to the event onset as a baseline for N1 calculation masks such non-spontaneous dynamics which more likely have functional significance. Accordingly, shorter time interval between sound-triggering button presses, have been reported to lead to smaller LRP amplitude (Hackley & Miller, 1995), and were associated with smaller N1 difference between self- and externally-generated sounds (Sanmiguel, Todd, & Schröger, 2013). Therefore, we believe it is preferable to use a baseline during which activity across different experimental conditions is similar and not inherently different due to the experimental conditions being compared.

In the current study, the corrected N1 amplitude evoked by self-generated sounds was enhanced relative to identical sounds passively heard. This was true using both baselines (−2600 to −2500 ms and −100 to 0 ms prior to button press). This is opposite to the attenuated N1 amplitude for self- vs. externally-generated sounds reported in previous studies (see Horvath, 2015 for a review). This might be explained by the fact that in our study, the evoked EEG activity in the motor-only condition was positive and much larger compared with that reported in previous studies. The correction we applied involved substracting the evoked activity in the motor-only condition from the evoked activity in the motor+sound condition. Thus, following substraction, the N1 amplitude in the corrected motor+sound condition became even more negative.

It should be noted that most studies that assess the auditory-evoked N1 component use headphones and not free-field speakers, as we did in the current study. When sounds are delivered through speakers, head movements can affect the relative position of the ears to the sound source, and therefore affect auditory processing and the evoked N1 amplitude. In the current study subjects were instructed to fixate on a display screen, thus stabilizing their head position throughout the experiment. Altough we cannot rule out differences in head position relative to sound source across conditions, we believe they are unlikely to explain the reported pattern of results. Aditionally, although we controlled for the auditory expectancy effect during self-generated sounds, some differences in level of temporal expectation between motor+sound and sound-only condition may still be present and cannot be ruled out (see Lange, 2013 for a review). In the current study, we found that despite non-significant differences in the distribution of IPIs, mean IPI across subjects was ~150 ms longer in the motor+sound compared with motor-only conditions. It can be argued that such differences in IPI could contribute to the differences we find in RP between motor+sound and motor-only conditions. Howver, the reported temporal differences in RPs greatly extend this difference in IPIs (starting ~1.4 seconds prior to button press). Moreover, using a similar experimental design, Ford and colleagues report differences in IPIs between silent and sound-triggering actions that were in the opposite direction, yet the differences in RPs between conditions were consistent with our findings (Ford et al., 2014). Therefore, it seems unlikely that differences in IPI can explain the differences in RPs between motor+sound and motor-only conditions reported in the current study.

According to the ideomotor theory (Greenwald, 1970), self-generated motor actions represent internalized mental images of their predicted sensory consequences. Our results support this view by showing that motor-related preparatory brain activity (RP) is different between sound-triggering and silent button presses. Thus, our results support this theory by showing that RP does not reflect motor planning only, but also involves early coding of the internal sensory image. Moreover, the ideomotor theory suggests that voluntary actions should lead to modulated sensory representation of the evoked reafferent stimulus (see Shin, Proctor, & Capaldi, 2010 for a review). This view is consistent with the current and previous physiological research showing modulated N1 amplitude evoked by self-generated sounds (Baess et al., 2011; J. Horvath, 2013; Sowman, Kuusik, & Johnson, 2012, Horvath, 2015) and behavioral findings showing that perceived loudness of self-generated sounds is modulated compared with sounds perceived in a passive manner (Reznik, Henkin, Levy, & Mukamel, 2015; Weiss, Herwig, & Schutz-Bosbach, 2011). It is still an open question how RP is modulated by the expectancy of action consequences with a probabilistic nature.

To conclude, in the current study we report that expectation of sensory action consequences modulates the temporal dynamics of neural activity preceding self-generated actions. We show that this modulation starts ~1.4 seconds prior to sound-triggering actions, suggesting the involvement of high-order motor areas in encoding the expected sensory consequences of self-generated actions. Taking into account such differences in temporal dynamics has important implications on the choice of baseline when comparing evoked responses to self vs. externally generated sounds.

## Acknowledgements

This study was supported by the I-CORE Program of the Planning and Budgeting Committee, and Israel Science Foundation Grant 51/11, Israel Science Foundation Grants 1771/13 and 2043/13, and Human Frontiers Science Project Career Development Award CDA00078/2011-C to R. M.; Sagol School of Neuroscience for D. R. and S. S and the Israeli Presidential Honorary Scholarship for Neuroscience Research to D. R. We thank laboratory members for fruitful discussion and comments on this manuscript.

## References

Alexander P., Schlegel, A., Sinnott-Armstrong, W., Roskies, A. L., Wheatley, T., & Tse, P. U. (2016). Readiness potentials driven by non-motoric processes. Consciousness and Cognition, 39, 38–47. doi:10.1016/j.concog.2015.11.011

Baess, P., Horvath, J., Jacobsen, T., & Schroger, E. (2011). Selective suppression of self-initiated sounds in an auditory stream: An ERP study. Psychophysiology, 48(9), 1276–1283. doi:10.1111/j.1469-8986.2011.01196.x

Benjamini, Y., & Hochberg, Y. (1995). Controlling the False Discovery Rate - a Practical and Powerful Approach to Multiple Testing. Journal of the Royal Statistical Society Series B-Methodological, 57(1), 289–300.

Blakemore, S. J., Smith, J., Steel, R., Johnstone, E. C., & Frith, C. D. (2000). The perception of self-produced sensory stimuli in patients with auditory hallucinations and passivity experiences: evidence for a breakdown in self-monitoring. Psychological Medicine, 30(5), 1131–1139. doi:10.1017/s0033291799002676

Brandi, M. L., Wohlschlager, A., Sorg, C., & Hermsdorfer, J. (2014). The Neural Correlates of Planning and Executing Actual Tool Use. Journal of Neuroscience, 34(39), 13183–13194. doi:10.1523/Jneurosci.0597-14.2014

Bulea, T. C., Prasad, S., Kilicarslan, A., & Contreras-Vidal, J. L. (2014).Sitting and standing intention can be decoded from scalp EEG recorded prior to movement execution. Frontiers in Neuroscience, 8. doi:10.3389/fnins.2014.00376

Christensen, M. S., Lundbye-Jensen, J., Geertsen, S. S., Petersen, T. H., Paulson, O. B., & Nielsen, J. B. (2007). Premotor cortex modulates somatosensory cortex during voluntary movements without proprioceptive feedback. Nature Neuroscience, 10(4), 417–419. doi:10.1038/nn1873

Cisek, P., & Kalaska, J. F. (2004). Neural correlates of mental rehearsal in dorsal premotor cortex. Nature, 431(7011), 993–996. doi:10.1038/nature03005

Crapse, T. B., & Sommer, M. A. (2008). Corollary discharge across the animal kingdom. Nature Reviews Neuroscience, 9(8), 587–600. doi:10.1038/nrn2457

Cui, F., Arnstein, D., Thomas, R. M., Maurits, N. M., Keysers, C., & Gazzola, V. (2014). Functional Magnetic Resonance Imaging Connectivity Analyses Reveal Efference-Copy to Primary Somatosensory Area, BA2. Plos One, 9(1). doi:10.1371/journal.pone.0084367

Cunnington, R., Windischberger, C., Deecke, L., & Moser, E. (2003). The preparation and readiness for voluntary movement: a high-field event-related fMRI study of the Bereitschafts-BOLD response. Neuroimage, 20(1), 404–412. doi:10.1016/s1053-8119(03)00291-x

De Jong, R., Wierda, M., Mulder, G., & Mulder, L. J. M. (1988). Use of partial stimulus information in response processing. Journal of Experimental Psychology: Human Perception and Performance, 14, 682–692.

Eimer, M. (1998). The lateralized readiness potential as an on-line measure of central response activation processes. Behavior Research Methods Instruments & Computers, 30(1), 146–156. doi:10.3758/Bf03209424

Eisenberg, M., Shmuelof, L., Vaadia, E., & Zohary, E. (2011). The Representation of Visual and Motor Aspects of Reaching Movements in the Human Motor Cortex. Journal of Neuroscience, 31(34), 12377–12384. doi:10.1523/Jneurosci.0824-11.2011

Eliades, S. J., & Wang, X. Q. (2003). Sensory-motor interaction in the primate auditory cortex during self-initiated vocalizations. Journal of Neurophysiology, 89(4), 2194–2207. doi:10.1152/jn.00627.2002

Ford, J. M., Palzes, V. A., Roach, B. J., & Mathalon, D. H. (2014). Did I Do That? Abnormal Predictive Processes in Schizophrenia When Button Pressing to Deliver a Tone. Schizophrenia Bulletin, 40(4), 804–812. doi:10.1093/schbul/sbt072

Ford, J. M., Roach, B. J., Faustman, W. O., & Mathalon, D. H. (2007). Synch before you speak: Auditory hallucinations in schizophrenia. American Journal of Psychiatry, 164(3), 458–466. doi:10.1176/appi.ajp.164.3.458

Ford, J. M., Roach, B. J., Faustman, W. O., & Mathalon, D. H. (2008). Out-of-synch and out-of-sorts: Dysfunction of motor-sensory communication in schizophrenia. Biological Psychiatry, 63(8), 736–743. doi:10.1016/j.biopsych.2007.09.013

Frank, M. (2001). Defective Recognition of One’s Own Actions in Patients With Schizophrenia. American Journal of Psychiatry, 158(3): 454–459

Fried, I., Mukamel, R., & Kreiman, G. (2011). Internally Generated Preactivation of Single Neurons in Human Medial Frontal Cortex Predicts Volition. Neuron, 69(3), 548–562. doi:10.1016/j.neuron.2010.11.045

Frith, C. D., Blakemore, S. J., & Wolpert, D. M. (2000). Explaining the symptoms of schizophrenia: Abnormalities in the awareness of action. Brain Research Reviews, 31(2–3), 357–363. doi:10.1016/s0165-0173(99)00052-1

Gallivan, J. P., McLean, D. A., Smith, F. W., & Culham, J. C. (2011). Decoding Effector-Dependent and Effector-Independent Movement Intentions from Human Parieto-Frontal Brain Activity. Journal of Neuroscience, 31(47), 17149–17168. doi:10.1523/jneurosci.1058-11.2011

Gratton, G., Coles, M. G., Sirevaag, E. J., Eriksen, C. W., & Donchin, E. (1988). Pre- and poststimulus activation of response channels: a psychophysiological analysis. Journal of Experimental Psychology: Human Perception and Performance, 14(3), 331–344.

Greenwald, A. G. (1970). Sensory feedback mechanisms in performance control: with special reference to the ideo-motor mechanism. Psychol. Rev., 7, 73–99.

Hackley, S. A., & Miller, J. (1995). Response Complexity and Precue Interval Effects on the Lateralized Readiness Potential. Psychophysiology, 32(3), 230–241. doi:10.1111/j.1469-8986.1995.tb02952.x

Haggard, P. (2008). Human volition: towards a neuroscience of will. Nature Reviews Neuroscience, 9(12), 934–946. doi:10.1038/nrn2497

Haggard, P., Clark, S., & Kalogeras, J. (2002). Voluntary action and conscious awareness. Nature Neuroscience, 5(4), 382–385. doi:10.1038/nn827

Haggard, P., & Eimer, M. (1999). On the relation between brain potentials and the awareness of voluntary movements. Experimental Brain Research, 126(1), 128–133. doi:10.1007/s002210050722

Haggard, P., & Whitford, B. (2004). Supplementary motor area provides an efferent signal for sensory suppression. Brain Research Cognitive Brain Research, 19(1), 52–58. doi:10.1016/j.cogbrainres.2003.10.018

Horvath, J. (2015). Action-related auditory ERP attenuation: Paradigms and hypotheses. Brain Research, 1626, 54–65. doi:10.1016/j.brainres.2015.03.038

Horvath, J. (2013). Attenuation of auditory ERPs to action-sound coincidences is not explained by voluntary allocation of attention. Psychophysiology, 50(3), 266–273. doi:10.1111/Psyp.12009

Horvath, J., Maess, B., Baess, P., & Toth, A. (2012). Action-Sound Coincidences Suppress Evoked Responses of the Human Auditory Cortex in EEG and MEG. Journal of Cognitive Neuroscience, 24(9), 1919–1931. doi:10.1162/jocn_a_00215

Hughes, G., Desantis, A., & Waszak, F. (2012). Mechanisms of intentional binding and sensory attenuation: The role of temporal prediction, temporal control, identity prediction, and motor prediction. Psychological Bulletin, 139, 133–151. doi:10.1037/a0028566

Hughes, G., Desantis, A., & Waszak, F. (2013). Attenuation of auditory N1 results from identity-specific action-effect prediction. European Journal of Neuroscience, 37(7), 1152–1158. doi:10.1111/ejn.12120

Jo, H. G., Wittmann, M., Hinterberger, T., & Schmidt, S. (2014). The readiness potential reflects intentional binding. Frontiers in Human Neuroscience, 8. doi:10.3389/Fnhum.2014.00421

Kornhuber, H. H., & Deecke, L. (1990). Readiness for Movement - the Bereitschaftspotential Story. Current Contents/Life Sciences 33(4), 14–14.

Kutas, M., & Donchin, E. (1980). Preparation to respond as manifested by movement-related brain potentials. Brain Res, 202, 95–115.

Lang, W., Cheyne, D., Kristeva, R., Beisteiner, R., Lindinger, G., & Deecke, L. (1991). Three-dimensional localization of SMA activity preceding voluntary movement. A study of electric and magnetic fields in a patient with infarction of the right supplementary motor area. Experimental Brain Research, 87(3), 688–695.

Lange, K.(2013).The ups and downs of temporal orienting: a review of auditory temporal orienting studies and a model associating the heterogeneous findings on the auditory N1 with opposite effects of attention and prediction. Frontiers in Human Neuroscience,7. doi:Artn26310.3389/Fnhum.2013.00263

Libet, B., Gleason, C. A., Wright, E. W., & Pearl, D. K. (1983). Time of conscious intention to act in relation to onset of cerebral activity (readiness-potential). Brain, 106(3), 623–642.

Makoshi, Z., Kroliczak, G., & van Donkelaar, P. (2011). Human Supplementary Motor Area Contribution to Predictive Motor Planning. Journal of Motor Behavior, 43(4), 303–309. doi:10.1080/00222895.2011.584085

Nicolelis, M. A., Baccala, L. A., Lin, R. C., & Chapin, J. K. (1995). Sensorimotor encoding by synchronous neural ensemble activity at multiple levels of the somatosensory system. Science, 268(5215), 1353–1358. doi:10.1126/science.7761855

Passingham, R. E. (1987). Two cortical systems for directing movement. Ciba Foundation Symposium, 132, 151–164.

Perez, O., Mukamel, R., Tankus, A., Rosenblatt, J. D., Yeshurun, Y., & Fried, I. (2015). Preconscious Prediction of a Driver’s Decision Using Intracranial Recordings. Journal of Cognitive Neuroscience, 27(8), 1492–1502. doi:10.1162/jocn_a_00799

Pedersen, J. R., Johannsen, P., Bak, C. K., Kofoed, B., Saermark, K., & Gjedde, A. (1998). Origin of human motor readiness field linked to left middle frontal gyrus by MEG and PET. Neuroimage, 8(2), 214–220. doi:10.1006/nimg.1998.0362

Quinn, B. T., Carlson, C., Doyle, W., Cash, S. S., Devinsky, O., Spence, C., Thesen, T., (2014). Intracranial Cortical Responses during Visual-Tactile Integration in Humans. Journal of Neuroscience, 34(1), 171–181. doi:10.1523/Jneurosci.0532-13.2014

Reznik, D., Henkin, Y., Levy, O., & Mukamel, R. (2015). Perceived Loudness of Self-Generated Sounds Is Differentially Modified by Expected Sound Intensity. PLoS One, 10(5). doi:ARTNe012765110.1371/journal.pone.0127651

Reznik, D., Ossmy, O., & Mukamel, R. (2015). Enhanced Auditory Evoked Activity to Self-Generated Sounds Is Mediated by Primary and Supplementary Motor Cortices. Journal of Neuroscience, 35(5), 2173–2180. doi:10.1523/Jneurosci.3723-14.2015

Sanmiguel, I., Todd, J., & Schröger, E. (2013). Sensory suppression effects to self initiated sounds reflect the attenuation of the unspecific N1 component of the auditory ERP. Psychophysiology, 50, 334–343. doi:10.1111/psyp.12024

Schurger, A., Sitt, J. D., & Dehaene, S. (2012). An accumulator model for spontaneous neural activity prior to self-initiated movement. Proceedings of the National Academy of Sciences of the United States of America, 109(42). doi:10.1073/pnas.1210467109

Shibasaki, H., & Hallett, M. (2006). What is the Bereitschaftspotential? Clinical Neurophysiology, 117(11), 2341–2356. doi:10.1016/j.clinph.2006.04.025

Soon, C. S., Brass, M., Heinze, H. J., & Haynes, J. D. (2008). Unconscious determinants of free decisions in the human brain. Nature Neuroscience, 11(5), 543–545. doi:10.1038/nn.2112

Sowman, P. F., Kuusik, A., & Johnson, B. W. (2012). Self-initiation and temporal cueing of monaural tones reduce the auditory N1 and P2. Experimental Brain Research, 222(1–2), 149–157. doi:10.1007/s00221-012-3204-7

Shin, Y. K., Proctor, R. W., & Capaldi, E. J. (2010). A Review of Contemporary Ideomotor Theory. Psychological Bulletin, 136(6), 943–974. doi:10.1037/a0020541

Smid, H. G. O. M., Mulder, G., & Mulder, L. J. M. (1987). The continuous flow model revisited: Perceptual and central motor aspects., Current trends in event-related potential research EEG supplement (Vol. 40, pp. 270–278). Amsterdam: Elsevier.

von Holst, E. (1954). Relations between the central Nervous System and the peripheral organs. Animal Behavior, 2(3), 89–94. doi:10.1016/S0950-5601(54)80044-X

Voss, M., Ingram, J. N., Haggard, P., & Wolpert, D. M. (2006). Sensorimotor attenuation by central motor command signals in the absence of movement. Nature Neuroscience, 9(1), 26–27. doi:10.1038/nn1592

Waters, F., Woodward, T., Allen, P., Aleman, A., & Sommers, I. (2012). Self-recognition Deficits in Schizophrenia Patients With Auditory Hallucinations: A Meta-analysis of the Literature. Schizophrenia Bulletin, 38(4), 741–750. doi:10.1093/schbul/sbq144

Weiss, C., Herwig, A., & Schutz-Bosbach, S. (2011). The self in action effects: selective attenuation of self-generated sounds. Cognition, 121(2), 207–218. doi:10.1016/j.cognition.2011.06.011

Wolpert, D. M., Ghahramani, Z., & Jordan, M. I. (1995). An internal model for sensorimotor integration. Science, 269(5232), 1880–1882. doi:10.1126/science.7569931

Yazawa, S., Ikeda, A., Kunieda, T., Ohara, S., Mima, T., Nagamine, T., Shibasaki, H., (2000). Human presupplementary motor area is active before voluntary movement: subdural recording of Bereitschaftspotential from medial frontal cortex. Experimental Brain Research, 131(2), 165–177. doi:10.1007/s002219900311

